# A natural marmoset model of genetic generalized epilepsy

**DOI:** 10.1101/2021.07.07.451379

**Authors:** Xiangyu Yang, Zhitang Chen, Ziying Wang, Guang He, Zhiqiang Li, Yongyong Shi, Neng Gong, Binglei Zhao, Eiki Takahashi, Weidong Li

## Abstract

As a common neurological disease, epilepsy has been extensively studied. Efforts have been made on rodent and other animal models to reveal the pathogenic mechanisms of epilepsy and develop new drugs as treatment. However, the features of current epilepsy models cannot fully mimic different kinds of epilepsy in human, asking for non-human primates models of epilepsy. The common marmoset (*Callithrix jacchus*) is a New World monkey that is widely used to study brain function. Here, we show a natural marmoset model of generalized epilepsy. In this unique marmoset family, generalized epilepsy was successfully induced by handling operation in some individuals. We mapped the marmoset family with handling-sensitive epilepsy and found that epileptic marmoset had an autosomal dominant genetic predisposition. Those marmosets were more sensitive to epilepsy inducers pentylenetetrazol (PTZ). By electrocorticogram (ECoG) recording, we detected epileptic discharge in marmoset with history of seizures. However, there was no significant change in the overall structure of epileptic marmoset brain. In summary, we report a family of marmosets with generalized seizures induced by handling operation. This epileptic marmoset family provides insights to better understand the mechanism of generalized epilepsy and helps to develop new therapeutic methods.

**Short summary:** Despite the rodent and other animal models on epilepsy, a common neurological disease, the understanding of the pathogenic mechanisms of epilepsy and treatment are yet to be advanced., Non-human primates are good experimental animals for modelling neurological disease. Here we report a common marmoset family with generalized epilepsy induced by handling stimulation. We mapped the epileptic marmoset family and found that epilepsy had a genetic predisposition. The present report deals in detail with the phenotypes and characteristics of epileptic marmosets. This natural marmoset reflect epilepsy model will help to better understand the mechanism of epilepsy and develop new therapeutic methods.

## Introduction

Epilepsy is a common chronic brain disorder characterized by recurrent seizures, affecting over 70 million people worldwide^[1, 2]^. A third of patients are idiopathic generalized epilepsies which exhibit a polygenic and heritable etiology^[3]^. In the past few decades, animals such as rodents have been used to investigate the mechanism and treatments of epilepsy ^[4]^. Due to the differences in genetic constitution and brain structure, rodent models can not fully mimic the human epilepsy. To address this issue, non-human primate models are considered as suitable models of nervous system disease because the non-human primate brain have very similar genetic, neurochemical, neurophysiologic, and structure features with human brains ^[5]^. Killam et al. firstly described an non-human primate baboon model of photosensitive epilepsy in 1966, which was characterized by intermittent light stimulation (ILS) induced seizures^[6]^. This was the first natural epileptic non-human primate model. However, so far no natural model of epilepsy in marmosets have been reported.

The common marmoset (*Callithrix jacchus*) is a small new world monkey, which has been frequently used because of its genetic constitution, body size, and other unique biological characteristics^[7, 8]^. Genome-wide data has been obtained from common marmosets and the results showed that most of the genes between marmosets and humans are highly conserved^[9, 10]^. This suggests that marmoset is a valuable biomedical model of primates. In epilepsy research, marmosets are mainly used for drug-induced epilepsy models and reliable for evaluation on antiepileptic drugs ^[11–13]^.

In this study, we showed a marmoset with generalized epilepsy for the first time. In this marmoset family, some individuals showed significant seizure phenotypes in response to handling stimulation. The seizure symptoms include limb convulsion, movement disorder, vomiting and salivation, which are typical in human epileptic seizures. Moreover, we found that this phenotype was stably inherited by the next generations. The phenotypes of marmoset were further identified through behavioral observations and in vivo ECoG recordings. Epileptic marmosets show more serious symptoms to PTZ treatment. We suggest this natural epileptic marmoset as a potential non-human primate model for understanding the mechanism of epilepsy.

## Methods

### Animals

All animal experiments were approved by Institutional Animal Care and Use Committee (IACUC) of Shanghai Jiao Tong University (No.10645, Shanghai, China) and the Animal Experiments Committee of CLEA (No.55-019CJ, CLEA Japan Inc., Tokyo, Japan). Twenty-one common marmosets (Callithrix jacchus, 45-118 months old, 280-400g) were reared at the CLEA Marmoset Breeding Facility (Gifu, Japan), and maintained on a 12-h light-dark cycle at 27°C and 50% humidity. Marmosets were allowed ad libitum access to water and food pellets(CMS-1M; CLEA Japan Inc.) with added vitamins C and D, calcium, and acidophilus. Hot water and comb honey were also added to soften the pellets and improve the animals’ preference for the food.

### Electrode implant surgery

For Invasive ECoG recording, marmosets were subcutaneously implanted with eight channel electrode^[14]^. Briefly, marmosets’ scalps were shaved to expose the skull under deep anesthesia condition. And dental drill was used to make hole on each implanted site. Then stainless miniature electrode (Jiangsu Braintech, China) were carefully inserted into the skull. Eight ECoG electrodes were separately implanted into the frontal cortex (AP +9.0mm, ML ±5mm), motor and premotor cortex (AP +4.0mm, ML ±5mm), parietal cortex (AP−3.0mm, ML ±5mm) and occipital cortex (AP-10mm, ML ±5mm). The electrodes were affixed to the skull with dental cement. After electrodes implantation, a protective cap baseplate was fixed on the skull using dental cement and electrode connector was covered with protective cap. Then antibiotics were intraperitonelly injected to prevent infection. Marmosets were allowed to recover from surgery for 2 weeks.

### ECoG data collection and analysis in free roaming marmoset

The marmosets were allowed to free roaming in the recording cage. A 8-channel neural recording device (NeuroAir 1.0, Jiangsu BrainTech, China) was connected with the electrode. The protection cap was remounted on the cap base. ECoG signals were recorded during the light phase of the cycle. The raw data was down-sampled at 500Hz and a notch filter with 60Hz was used to remove power frequency interference during data acquisition. According to the non-stationary characteristics of ECoG signals, data of the same duration is selected for spike-waves detection and number counting.

### PTZ susceptibility testing

The pentylenetetrazol (S4587; Selleck Chemicals, USA) test was conducted based on previous study during the light phase of the cycle^[12]^. Each marmoset was habituated in an individual observation cage for at least 30 min prior to PTZ test. To assess the susceptibility of PTZ triggered seizure, the marmosets were injected with 35mg/kg PTZ intraperitoneally. Before PTZ treatment, their natural behaviors were monitored for 60 minutes. After PTZ injection, each marmoset was immediately placed into the observation cage. Seizures of marmosets were recorded for 60 minutes and evaluated through the revised Racine’s seizure scale.

### Seizure classification

The seizure behaviors were evaluated based on revised Racine’s seizure scale for marmoset proposed in previous study ^[12, 15]^. The seizure activities of marmosets were classified as follows:

I. Mouth cleaning-like behavior (rubbing the face along the perch and wire mesh);
II. Head clonus/shaking (rapid and violent movement of the head);
III. Forelimb clonus;
IV. Bilateral forelimb clonus, straub tail, postural impairment;
V. Generalized clonic seizures.

### Behavior analysis

Behaviors were recorded using a video camera. The locomotion events were recorded by movements of the animal involving both limbs, such as walking, jumping, running, and climbing. These events were counted in 5 minutes intervals for 60 minutes; The whole process was divided into two phases: the phase I (0-10 min) and phase II (11-60 min). We also recorded the time spent on scratching and counted the events of head shakes/Mouth cleaning.

### Morphological observation (In situ hybridization)

The brains asymptomatic marmosets (n=3) and epileptic marmosets (n=4)) were dissected after perfusion, fixed in Tissue Fixative (Gonostaff, Tokyo, Japan), embedded in paraffin, sectioned (6-μm-thick slices) and stained according to previous report using beta-actin probe (Gonostaff, Tokyo, Japan)^[16]^.

### Statistical analysis

Data were analyzed using GraphPad Prism software ver.6 (GraphPad, CA, USA). In all experiments, the experimenters were blind to the group and treatment of the marmosets. The data were analyzed with Student’s t-tests or two-way ANOVA, followed by Fisher's LSD tests for multiple group comparisions. Values were presented as means ± standard errors of the means (SEM). Results with P < 0.05 were regarded as statistically significant.

## Results

### Discovery of marmoset with generalized epilepsy

Epileptic marmosets were discovered as a result of routine handling for health check such as weight measurement. Some of the marmosets showed significant epileptic seizures during handling. By tracing the seizure history in this family, the first epileptic marmoset was identified born in 2002. This family, which has ancestry from South American, has 52 marmosets in four generations. The incidence rate of epileptic seizures in four generations are 2/2(G I), 15/22(G II), 8/16(G III), 4/12(G IV), respectively (Fig 1A). With mapping the marmoset family, it indicated that this handling-induced seizure phenotype was stably inherited and had an autosomal dominant genetic predisposition.

**Fig 1.**
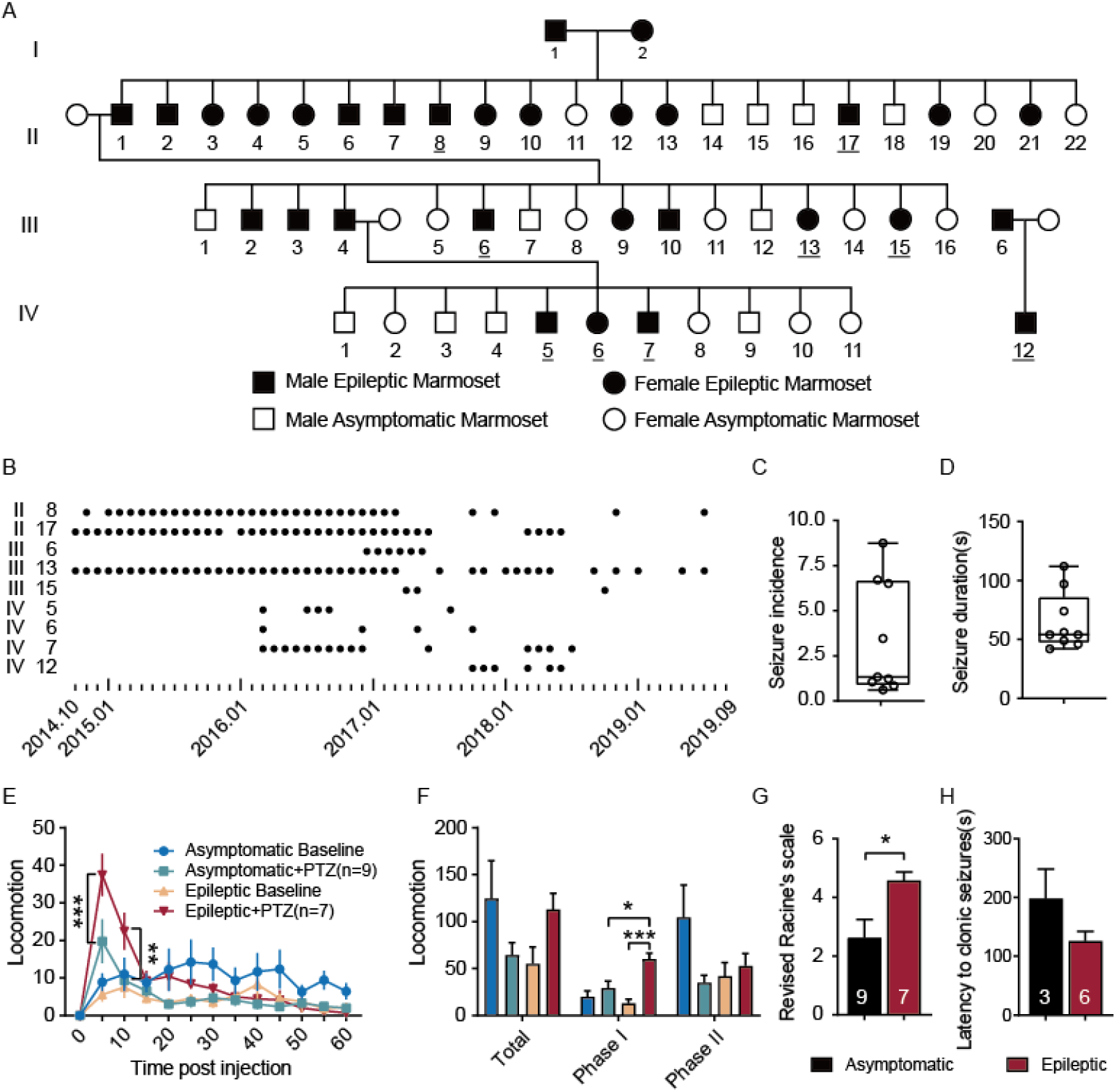
Marmoset family with generalized epilepsy. A. Pedigree of the epileptic marmoset family. B. Seizure records of the samples involved in the study. C. The seizure incidence of epileptic marmosets (n=9). D. The seizure duration statistical result of handling induced epilepsy with video records (n=9). E. Time course of locomotion after PTZ injection. F. Total duration of locomotion during the phase I (0–10 min) and phase II (11–60 min). G. Seizure scores in marmosets treated with PTZ. H. Latency to clonic (IV/V) seizures of PTZ-treated marmosets. Data are mean ± SEM. Scale bar: 5mm. \**P* <0 .05; \*\**P* <0 .01; \*\*\**P* <0 .001;

Seizures were triggered by handling in these marmosets. The seizure started with the clonic fore limb followed by crows myoclonia and generalized Tonic-Clonic. Since October 2014, seizures were recorded during the monthly body weight measurement in nine alive marmosets. We found that the earliest onset of seizure in the marmoset was about 11 months old (Sup. Table 1). Different epileptic marmosets had variable seizure incidences. Not every handling can induce seizures. Epileptic seizures records during the observation period were shown in Fig 1B. Approximately 3.38 seizures could be induced per 12 handling (Fig 1C, n = 9, 3.38 ± 1.04). Each seizure lasted about 64.89 seconds (Fig 1D, n = 9, 64.89 ± 8.15). We also tested light sensitivity in epileptic marmosets and found that different frequencies of light stimulation did not induce seizures.

PTZ-induced seizure susceptibility was assessed in marmosets. The epileptic marmoset exhibited more severe symptoms (Fig 1E-F). After PTZ treatment, the locomotion events increased in epileptic marmosets, while it decreased in asymptomatic marmosets. PTZ administration also induced significant changes in other behaviors, such as scratching, head shakes/Mouth cleaning behaviors(Sup. Table 2 and Sup. Fig 1). Epileptic marmosets also showed higher Racine’s score compared with asymptomatic marmosets (Fig 1G.*P* = 0.019). In PTZ test, six-sevenths of epileptic marmosets showed clonic (IV/V) seizures, and three ninths of asymptomatic marmosets developed clonic seizures. In epileptic marmoset group the latency of PTZ-induced clonic (IV/V) seizures were shorter(Fig 1H. *P* = 0.12). Those results indicated that epileptic marmosets have higher PTZ sensitivity.

### Characterization of brain activity and structure in epileptic marmosets

The ECoG recording pattern is shown in Fig. 2A. During handling for recording, epileptiform whole-brain discharge was observed in all three epileptic marmosets, with the frequency focused at 1-6Hz (Fig 2B-E). In the ECoG data from free roaming epileptic marmosets (10 hours), we also detected epileptic spikes using instantaneous envelope model(Fig 2F). However, in videos analysis, we did not observe obvious seizure behavior. By contrast, there were fewer epileptic spike waves in the asymptomatic marmosets than ...(Fig 2G, Asy:5.25±2.49, Epi:739 ± 305.5, *P* = 0.035).

**Fig2.**
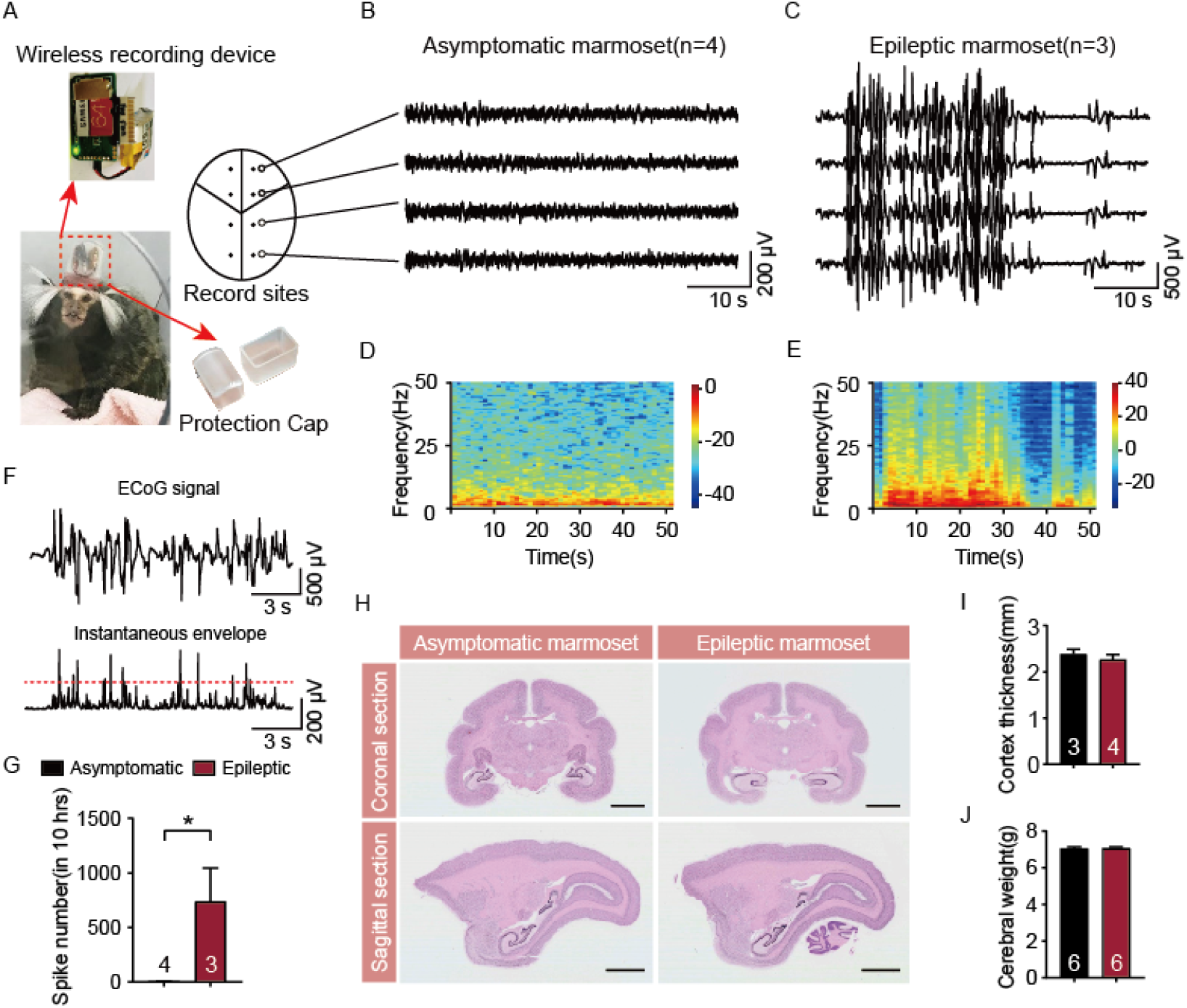
Phenotypic analysis of epileptic marmosets. A. Schematic diagram of EEG recording in marmosets. B,C. Typical ECoG traces after handling in asymptomatic marmosets and epileptic marmosets. D,E. Spectrogram corresponding to typical ECoG trace in asymptomatic marmosets and epileptic marmosets. F. Spike detect modelling. G. Number of spikes during 10 hours free-roaming. H. Image of Beta-actin in situ hybridization for brain structure comparison. I. Cortical thickness measurements in asymptomatic and epileptic marmosets. J. Cerebral weight measurements in asymptomatic and epileptic marmosets. Data are mean ± SEM. Scale bar: 5mm. \**P* <0 .05. (T-test)

Preliminary actin in situ hybridization of epileptic marmoset brains showed no significant structural differences (Fig 2H). No obvious changes were observed with cortex thickness and cerebral volume between asymptomatic and epileptic marmosets.

## Discussion

In this study, we showed a family of reflect epileptic marmosets with genetic predisposition. After a long period of retrospective investigation and observation, we tagged epilepic individuals in this family and mapped the family pedigree, and specified the behavioral characteristics of epileptic marmosets.

Autosomal dominant inheritance with reduced penetrance was observed in this marmoset family with handling sensitivity. Unlike photosensitive baboons, epileptic marmosets are not sensitive to different frequencies of light. Since the epileptic marmosets reported in other studies have similar behavioral phenotypes, researchers attribute it to a viral infection^[5]^. Based on our observations, a genetic etiology may be the causes of the epileptic seizure in this marmosets family. Hence, future studies are needed to reveal the cause of epileptic phenotypes, especially by using genetic analysis method. We chose whole-genome sequencing to screen the candidate pathogenic genes. But so far, no specific mutation has been found in this family.

In vivo electrophysiological results showed that handling induced significant epileptic seizures, but lacked synchronous behavior phenotypes. This suggests that a more complex mechanism could be underlying behavioral seizures. Due to the limitation of ECoG recording, we were unable to record the electrical activity of the deep brain regions, which limits our exploration of the pathogenesis of seizures. At present, we have observed significant seizure discharges after handling, but more elaborate recordings are needed to reveal the characteristics and mechanisms of seizure origin in marmosets brain. PTZ susceptibility testing showed that epileptic marmosets produced more pronounced epileptic phenotypes, suggesting the existence of neurological functional variations. However, preliminary structural observations showed no significant abnormalities in the brain structure of epileptic marmosets, suggesting that this neurological abnormality exists at a more detailed structural level and requires more sophisticated imaging of neural circuits and synaptic structures.

Combined with the above results, the discovery of this familial generalized epileptic marmosets will further enrich the application of non-human primate animals in the field of epilepsy research. Meanwhile, this epileptic marmoset model is another natural epilepsy model but different from photosensitive baboons, which is conducive to the study of the mechanism and treatment of epileptic seizures. In addition, further genetic research will clarify the genetic mechanism of this familial generalized epilepsy, which could help advance the pathological studies in clinical.

## ACKNOWLEDGMENTS

This work was supported by the National Key Research and Development Program of China (2018YFE0126700), the National Natural Science Foundation of China (82001372) and Shanghai Municipal Commission of Science and Technology Program (21dz2210100).

## References

1. Roland D Thijs, Rainer Surges, Terence J O’Brien, and Josemir W Sander, Epilepsy in adults. The Lancet, 2019. 393(10172): p. 689–701.

2. Annamaria Vezzani, Jacqueline French, Tamas Bartfai, and Tallie Z Baram, The role of inflammation in epilepsy. Nature reviews neurology, 2011. 7(1): p. 31.

3. Pierre Jallon and Patrick Latour, Epidemiology of idiopathic generalized epilepsies. Epilepsia, 2005. 46: p. 10–14.

4. Brian P Grone and Scott C Baraban, Animal models in epilepsy research: legacies and new directions. Nature neuroscience, 2015. 18(3): p. 339–343.

5. Leah Croll, Charles A Szabo, Noha Abou‐Madi, and Orrin Devinsky, Epilepsy in nonhuman primates. Epilepsia, 2019. 60(8): p. 1526–1538.

6. KF Killam, R Naquet, and J Bert, Paroxysmal responses to intermittent light stimulation in a population of baboons (Papio papio). Epilepsia, 1966. 7(3): p. 215–219.

7. Hideyuki Okano, Erika Sasaki, Tetsuo Yamamori, Atsushi Iriki, Tomomi Shimogori, Yoko Yamaguchi, Kiyoto Kasai, and Atsushi Miyawaki, Brain/MINDS: a Japanese national brain project for marmoset neuroscience. Neuron, 2016. 92(3): p. 582–590.

8. Noriyuki Kishi, Kenya Sato, Erika Sasaki, and Hideyuki Okano, Common marmoset as a new model animal for neuroscience research and genome editing technology. Development, growth & differentiation, 2014. 56(1): p. 53–62.

9. The Marmoset Genome Sequencing, Kim C Worley, Wesley C Warren, Jeffrey Rogers, Devin Locke, Donna M Muzny, Elaine R Mardis, George M Weinstock, Suzette D Tardif, and Kjersti M Aagaard, The common marmoset genome provides insight into primate biology and evolution. Nature genetics, 2014. 46(8): p. 850.

10. Chentao Yang, Yang Zhou, Stephanie Marcus, Giulio Formenti, Lucie A Bergeron, Zhenzhen Song, Xupeng Bi, Juraj Bergman, Marjolaine Marie C Rousselle, and Chengran Zhou, Evolutionary and biomedical insights from a marmoset diploid genome assembly. Nature, 2021: p. 1–9.

11. Michael P Hill, Erwan Bezard, Steven G McGuire, Alan R Crossman, Jonathan M Brotchie, Ann Michel, Renee Grimée, and Henrik Klitgaard, Novel antiepileptic drug levetiracetam decreases dyskinesia elicited by L‐dopa and ropinirole in the MPTP‐lesioned marmoset. Movement disorders: official journal of the Movement Disorder Society, 2003. 18(11): p. 1301–1305.

12. João C Bachiega, Miriam M Blanco, Patrícia Perez-Mendes, Simone Maria Cinini, Luciene Covolan, and Luiz E Mello, Behavioral characterization of pentylenetetrazol-induced seizures in the marmoset. Epilepsy & Behavior, 2008. 13(1): p. 70–76.

13. Josy Carolina C Pontes, Thiago Z Lima, Claudio M Queiroz, Simone M Cinini, Miriam M Blanco, and Luiz E Mello, Seizures triggered by pentylenetetrazol in marmosets made chronically epileptic with pilocarpine show greater refractoriness to treatment. Epilepsy research, 2016. 126: p. 16–25.

14. Sabyasachi Roy and Xiaoqin Wang, Wireless multi-channel single unit recording in freely moving and vocalizing primates. Journal of neuroscience methods, 2012. 203(1): p. 28–40.

15. Akiyoshi Ishikawa, Yuri Mizuno, Keita Sakai, Takehiro Maki, Ryo Tanaka, Yasuhiro Oda, Kimie Niimi, and Eiki Takahashi, Kainic acid-induced seizures in the common marmoset. Biochemical and Biophysical Research Communications, 2020.

16. Yoshinori Shimamoto, Kimie Niimi, Hiroshi Kitamura, Sae Tsubakishita, and Eiki Takahashi, In situ hybridization study of CYP2D mRNA in the common marmoset brain. Experimental animals, 2016. 65(4): p. 465–471.

