## Supplementary materials for "A natural marmoset model of genetic generalized epilepsy"

**Supplementary Materials for**
**A natural marmoset model of genetic generalized epilepsy.**

### Materials and Methods

#### ECoG analysis,

ECoG data from 9 am to 7 pm were extracted for analysis, with a total of 10 hours. The raw data was down-sampled at 500 Hz and a notch filter with 60 Hz was used to remove power frequency interference during data acquisition. Data were baseline corrected by subtraction of the mean of all channels, re-referenced to the average of all channels and digitally filtered offline at 0.5-200 Hz.

#### Epileptic spike detection

The wavelet transform has multi-resolution characteristics, and a more accurate signal can be observed. By properly selecting the wavelet basis function, the wavelet transform can have the ability to characterize the local characteristics of the signal in both time and frequency domains. According to the non-stationary characteristics of ECoG signals, data of the same duration is selected for wavelet transform. Because the cross-correlation value of the db4 function and the epilepsy signal is the largest, db4 is selected as the wavelet basis function in this study, and a 7-level-decomposition transform is used to decompose the signal, namely the detail coefficients D1, D2, D3, D4, D5, D6, D7 and the approximate coefficient A7[1]. By setting a threshold to filter out maxima points caused by noise and other small interferences, the number of the coefficient of each layer is higher than the threshold, that is, the number of spikes is counted. Compare and analyze the spikes number of different phenotypes of marmosets.

#### Reference.

1. Tzimourta, K. D., Tzallas, A. T., Giannakeas, N., Astrakas, L. G., Tsalikakis, D. G., & Tsipouras, M. G.. *Epileptic seizures classification based on long-term EEG signal wavelet analysis. In International Conference on Biomedical and Health Informatics*. ICBHI, Singapore 2017, November. p. 165-169.

**Table 1** The detail information of epileptic marmosets in this unique family.

| Epileptic marmoset ID | DOB | Age（Months） | Gender | Start Recording | Seizure Start（month age） | Number of Seizures |
|---|---|---|---|---|---|---|
| **II 2** | 2013.05 | 78 | M | 2014.10 | - | 33 |
| **II 4** | 2011.09 | 98 | M | 2014.10 | - | 32 |
| **III 6** | 2013.06 | 77 | F | 2014.10 | - | 43 |
| **III 7** | 2013.11 | 69 | F | 2014.10 | - | 3 |
| **III 9** | 2009.05 | 126 | M | 2014.10 | - | 6 |
| **IV 7** | 2014.11 | 60 | M | 2014.12 | 2016.03(16) | 5 |
| **IV 8** | 2014.11 | 56 | F | 2014.12 | 2016.03(16) | 4 |
| **IV 9** | 2015.04 | 55 | M | 2015.05 | 2016.03(11) | 15 |
| **IV 12** | 2015.02 | 57 | M | 2015.03 | 2017.10(32) | 6 |

All information in this table was collected from Oct. 2014 to Sep. 2019.

**Table 2** The frequencies of various behaviors in marmosets treated with PTZ

| | **Asymptomatic marmoset(n=9)** | | **Epileptic marmoset(n=7)** | |
|---|---|---|---|---|
| | Baseline | PTZ | Baseline | PTZ |
| **Natural behaviors** | | | | |
| Locomotion | 124.3±40.5 | 64.2±13.5 | 59.7±17.9 | 112.7±17.3 |
| Scratching (s) | 32.7±8.1 | 6.1±2.3 | 19.3±3.4 | 3.6±0.5 |
| **Early convulsive behaviors** | | | | |
| Mouth cleaning (number of events) | 4.4±0.5 | 12.6±2.4 | 3.7±0.6 | 23.9±2.8 |
| Head clonus/shakes (number of events) | 4.7±0.6 | 10.6±1.2 | 4.9±0.8 | 18.1±1.6 |

Each animal was observed for 60 min both before (baseline) and after PTZ injection.

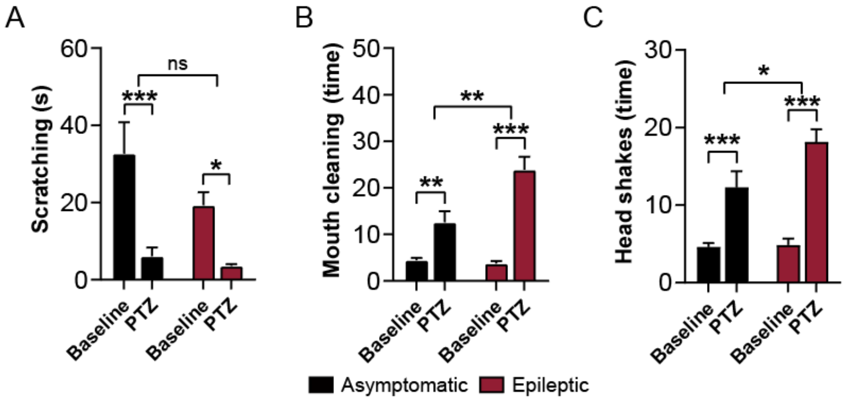


**Sup. Fig1.** Various behaviors in marmosets treated with PTZ. A.Scratching, B.Mouth cleaning, C. Head shakes bahaviors in asymatomatic and epileptic marmoset. Data are expressed as the means ± SEMs. Two-way ANOVA followed by Fisher's LSD test was used to compare behaviors between each group. *$P$ < 0.05, **$P$ < 0.01, ***$P$ < 0.001.
